# NetAllergen, a random forest model integrating MHC-II presentation propensity for improved allergenicity prediction

**DOI:** 10.1101/2022.09.22.509069

**Authors:** Yuchen Li, Peter Wad Sackett, Morten Nielsen, Carolina Barra

## Abstract

Allergy is a pathological immune reaction towards innocuous protein antigens. Although only a narrow fraction of plant or animal proteins induce allergy, atopic disorders affect millions of children and adults and cost billions in healthcare systems worldwide. In-silico predictors can aid in the development of more innocuous food sources. Previous allergenicity predictors used sequence similarity, common structural domains, and amino acid physicochemical features. However, these predictors strongly rely on sequence similarity to known allergens and fail to predict protein allergenicity accurately when similarity diminishes. In addition, ‘allergen’ is a broad terminology that may include different compounds, hindering the classification task. To overcome these limitations, we collected allergens from AllergenOnline, a curated database of IgE-inducing allergens, carefully removed allergen redundancy with a novel protein partitioning pipeline, and developed a new allergen prediction method, introducing MHC presentation propensity as a novel feature. NetAllergen outperformed a sequence similarity-based BLAST baseline approach, and previous allergenicity predictor AlgPred 2 when similarity to known allergens is limited. NetAllergen is available as a web service (services.healthtech.dtu.dk/service.php?NetAllergen-1.0) and can predict allergenicity from a protein sequence.

## INTRODUCTION

Currently, allergy affects as many as 20% of the worldwide population^1^, and only in the US, healthcare treatments cost several billions of dollars annually^2^. Moreover, the prevalence of allergic diseases has increased in the past decades, and it is likely to rise in the future^3^.

Allergy is characterised by a pathological immune reaction, triggered by normally ubiquitous and innocuous protein antigens in susceptible individuals^1^. In addition, allergy is a wide terminology that encompasses a number of diseases primarily sharing an IgE-dependent pathway^4^. Once the allergen contacts the skin or mucosal tissues, antigen-presenting cells (APCs) uptake and digest allergens into small peptides. Peptides with specific motifs are next presented by the major histocompatibility complex (MHC) class II molecules to activate type 2 T helper (Th2) cells. In susceptible individuals, Th2 cells further stimulate B cells to produce IgE antibodies in the first sensitization stage. An individual develops an allergy when the immune system is re-exposed to the allergen. The allergen is targeted by multiple IgE antibodies, which bind to Fc receptors in basophils and mast cells activating the degranulation process^4^. The reasons why only a small number of proteins induce allergy is largely unknown. However, it has been suggested that allergens share specific features that, according to the route and level of exposure, genetic predisposition, and environmental factors, can develop into allergies^5,6^.

MHC class II antigen presentation plays an essential role to mediate Th2 and IgE-producing immune responses in allergy development. Recently, genome-wide association studies (GWAS) have revealed that several allergies are strongly associated with specific HLA alleles, in particular for HLA-DQ and HLA-DRB^7^. For example, allergic rhinitis has shown a strong association with HLA-DRB1*04:01, HLA-DQB1*02:02, and HLA-DQB1*03:01^8^. HLA-DQA1*01:03-DQB1*06:01 was found to be associated with wheat allergy, and patients who had celiac diseases carried HLA-DQA1*05:01-DQB1*02:01 and HLA-DQA1*03:01-DQB1*03:02^9^. In allergic respiratory disease studies, HLA-DRB1*15:01 has been found to be associated with mite (Der p 1)^10^ and birch pollen (Bet v 1) allergens^11^. However, the same allele was negatively associated with gelatine^12^ and cow’s milk^13^. GWAS studies not only point to HLA class II specific allele associations but also suggest that specific amino acid positions in the peptide binding groove shared among different alleles provide susceptibility or protection against different types of allergies^8,14^. Together these findings strongly suggest that differential MHC-II presentation is involved in allergy.

Various methods have been utilized to identify common patterns in allergens and predict their allergenicity. Initially, relational databases were used to cluster allergens based on their protein sequence or shared structural domains, biochemical, and functional properties^5,15,16^. Both Blast and Allermatch, another sequence-comparing method, were used to find shared amino acid segments and search query sequences against allergen databases, respectively^17,18^. Later, motif-profile predictions improved prediction by searching for similarities in potential functional domains^19^. Machine learning-based methods including features such as known IgE epitopes, peptide motifs, and dipeptide compositions, have also been widely applied to predict allergenicity^20,21^. These include Allerdictor, which uses K-mer frequencies in a support vector machine (SVM)^22^, and AllergenFP based on Tanimoto coefficient similarity search^23^. Q-repeats, 3D structural similarities, and sequential similarities were incorporated into the decision tree in AllerCatPro^24^. The auto- and cross-covariance of amino acids in a sequence were adopted in AllerTOP v2^25,26^. AlgPred 2 predicts sequences with hybrid scores by integrating BLAST, and motif features using a Random Forest model^27^. However, to the best of our knowledge, no predictor has used MHC presentation in allergenicity prediction.

In this study, we developed a random forest predictor, NetAllergen, using MHC class II peptide presentation propensity for HLA-DR and HLA-DQ alleles in combination with physicochemical properties, structural information, and evolutionary information. We investigated and validated the relevance of MHC presentation propensity and other features for allergenicity predictions. NetAllergen was benchmarked against a sequence-based BLAST model and other state-of-the-art predictors and demonstrated to achieve higher performance, particularly for allergens that share limited similarity to previously known allergens.

## MATERIAL AND METHODS

### Training dataset

2,171 allergen protein IDs were retrieved from AllergenOnline, a curated database of IgE-producing allergens (On September 2020)^28^. Available FASTA sequences were downloaded from NCBI and filtered to include protein sequences that ranged in length from 50 to 1,000 (N=2082). To remove redundancy in the dataset, a Hobohm 1^29^ algorithm with 90% sequence identity was applied, resulting in a set of 686 non-redundant allergens covering 187 species. In short, the Hobohm 1 algorithm sorts the proteins by sequence length from longest to shortest. The first protein is automatically added to a ‘unique’ list, then the second protein is compared to the protein in the unique list. If the identity exceeds 90% the protein is removed, otherwise it is appended to the unique list. This process is iterated throughout the sorted protein list. Sequence identity was determined by pairwise alignment as the percentage of identical amino acids with regard to the query protein length.

For each allergen, non-allergen protein candidates were selected by searching for matching species names with the allergen and excluding results with the following keywords in NCBI: allergen, hypothetical, uncertain, predicted, probable and putative. Additionally, sequence lengths were limited to ±20% of the allergen length. Up to five candidates were randomly selected with BLASTP E-values (NCBI-Blast 2.12.0^30–32^) higher than 1×10^−10^ against the allergens (to avoid including sequences with high similarity to known allergens that could be untested antigens, Supp. Fig 1), and 1×10^−100^ against any sequence in the dataset (to avoid including repeated sequences). The number of non-allergens collected per allergen subset may vary (below five) because of the limited set of proteins for some species. The negative dataset consisted of 2,477 non-allergic protein sequences.

### Dataset homology reduction

In short, similar allergens were clustered together using BLAST E-score and minimum spanning tree (MST). First, the sequence similarity between each allergen in the dataset was determined by BLASTP using an all-against-all alignment. If the query did not provide a significant hit to a protein in the database, the E-score was set to a value of 10, scoring as non-aligned proteins. A pairwise comparison matrix of E-values was obtained to generate clusters by the MST algorithm, resulting in all sequences sharing similarity to at least one common neighbour being clustered together. All clusters when then randomly split in partitions to train the model. The training dataset was further reduced by different thresholds by removing the proteins that, after clustering with MST, had more than 20 neighbours/edges. The final positive dataset had 537 allergic proteins after removing homologous sequences. After clustering the allergens, all the non-allergens associated with each allergen (sharing the same species and length range) were grouped together. The clusters from MST were randomly distributed into five partitions with similar sizes.

### Feature constructions

#### MHC-II presentation propensity scores

NetMHCIIpan 4.0^33^ was used with a percentile rank below 10% to predict the number of peptides with a likelihood of being presented on MHC class II from a given protein in the dataset, using a sliding window of 15. From these numbers, two different features were collected. Binder 1 includes the counts for HLA-DRB1*04:01, HLA-DQA1*02:01-DQB1*02:02, HLA-DQA1*04:01-DQB1*03:01, HLA-DQA1*01:03-DQB1*06:01, HLA-DQA1*03:01-DQB1*03:02, and HLA-DQA1*05:01-DQB1*02:01 and binder 2 the counts for HLA-DRB1*15:01. These sets of alleles were selected as they were previously demonstrated to be positively (binder 1) or negatively/positively (binder 2) associated with allergies. Final MHC presentation propensity scores were calculated by summing the binders over different alleles normalised by sequence length.

#### Physicochemical properties

The physicochemical properties included average residue weight, average charge, and isoelectric point. Amino acids were also classified into tiny, small, aliphatic, aromatic, non-polar, polar, charged, basic, and acidic groups by size, structure, and charge. These were obtained using Pepstats from EMBL-EBI tools^34^A.dditionally, hydropathy indices were calculated using the Kyte-Doolittle scale^35^. A window size of 9 was used to determine position-specific hydropathy indices. Based on the sequential indices, we calculated an overall score for a protein by counting the number of hydrophobic or hydrophilic residues in a sliding window of 5. If all consecutive residues within a window had the same hydropathy tendency, a count of one was added to the overall score. To eliminate the influence of length on feature value, the scores were normalised by the sequence length.

#### Local structural features

Structural features were represented by alpha helix, beta sheet, relative solvent accessibility (RSA), and disordered regions. These features were predicted by using NetSurfP 3.0^36^. NetSurfP predicts residue-specific secondary structures. The occurrences of 5 alpha helix and 3 beta sheet residues within a sliding window of 7 summarised both alpha-helix and beta-sheet counts per protein. For RSA, the average value in the same window size was calculated, adding 1 if the value exceeded the 0.3 threshold. Additionally, under the same window size setting, the presence of more than three residues with a disorder higher than 0.5 was considered as 1 in the overall disorder. Likewise, these structural features were normalised by the sequence length.

#### Amino acid compositions and evolutionary information

The relative amino acid frequencies in the protein sequence were represented as 20 amino acid composition features (AAC). Further, the evolutionary information was represented by the position-specific scoring matrix (PSSM). PSI-BLAST was used to produce the PSSM^31^. The sequences in the training datasets were used as a BLAST database for searching. We replaced the PSSMs of query sequences with no hits with the default scoring matrix BLOSUM62 due to the database limit. We applied the autocovariance (AC) transformation to vectorize the PSSM for a protein^37^,

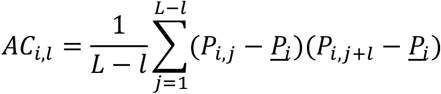

where *i* represents a specific amino acid, *l* is the lag to calculate the covariance, *L* is the protein sequence length, *j* is the amino acid position, *p*_*i,j*_ is a score of amino acid *i* at position *j* in the PSSM. As a result of vectorizing the PSSM, 20 autocovariance features are generated.

### Model development

Nested cross-validation was used for model training and evaluation. Based on the data partitioning, the outer layer had five-folds, and the inner layers had four folds to ensure that the partitions were mutually independent. Grid searches were used to determine hyperparameters and model selection were carried out on the training and validation datasets. A wide set of hyperparameters was tested including the number of estimators, the maximum depth of the decision tree, the minimum samples for split and leaf, and the impurity criterion (Supp. Table 1). The outer test sets were predicted using a mean of the ensemble of the selected optimal models.

On top of the partitioning and nested cross-validation, ten different random seeds were used to randomly distribute the clusters in the different partitions. Ten models were generated from the ten different seeds used for partitioning. The final model was made as an ensemble by taking the mean of ten different seed models.

All the random forest models were developed independently using this same pipeline. Performance evaluation was made on the entire data set by concatenating the predictions from the five outer test sets in the nested cross-validation setup. The input data were standardised to eliminate the effect of different feature scales using a z-score transformation. The z-scores were calculated based on the means and standard deviations of each training dataset. Data in validation and test datasets were normalised by the means and standard deviations from the training datasets used for model construction.

### Baseline model

Allergens in the training dataset were used to construct the BLAST database. Evaluation sequences were searched against the database and the E-values were extracted as the predictions of the baseline model.

### Evaluation datasets

Two independent evaluation datasets were collected. The first one was collected from the AlgPred 2 evaluation dataset^27^. Here, identical duplicates were removed leaving 765 allergen and 2,015 non-allergen sequences. The second was a collection of allergens retrieved from Allergen Nomenclature^38^, Allerbase^39^, COMPARE^40^, Allergome^41^, and SDAP^16^ databases (On December 2021). Allergenic protein sequences that were identical to positive sequences in the training dataset were excluded. Next, the same procedure to collect non-allergens as for the training dataset was applied. This evaluation dataset contained 1,065 positives and 4,914 negative proteins (Supp. Table 2).

### Evaluation metrics

The performance of each model was evaluated by AUC, AUC 0.1, and positive predictive value (PPV). AUC 0.1 was calculated as the area integrated to a 10% false positive rate. For PPV, predicted values were first sorted in descending order, next PPV was obtained from the proportion of true positive targets within the top N predictions, where N was the number of allergens.

In order to assess if the performance of the two models were significantly different, a p-value was calculated using bootstrapping. In this setup, a sample with replacement of the same size as the evaluation dataset was taken from the paired predictions from two models to determine whether the null hypothesis of the two models performing at par was rejected. This resampling was performed 10,000 times, and the p-value calculated by counting the occurrences of the AUC of one model was greater than that of another divided by the number of samplings.

## RESULTS

### Homology reduction pipeline

One of the big and current challenges in machine learning approaches is data leakage, which originates from data used for the model to learn, with high redundancy to what is later evaluated on^42^. This issue not only overestimates model accuracy but also curbs its generalisation and therefore hinders its prediction ability. In particular, the multiple allergen databases that exist contain a great number of homologous sequences or iso-allergens^39^. While this redundancy may facilitate the task of finding homologous candidate allergens in unexplored species, it might derange the effort to find new allergen families or sequence-unrelated unknown allergens. Hence, we first developed a pipeline to efficiently remove and cluster similar sequences from the allergens dataset using BLAST sequence alignment and minimum spanning tree (MST), which connects into single clusters all proteins sharing sequence similarity over a threshold.

Allergen protein sequences were collected from AllergenOnline^28^ and filtered to remove atypically short and long sequences, highly redundant or identical sequences (detailed in the Materials and Methods section, Fig 1A). After this, 686 allergen sequences were clustered by MST using all-against-all BLAST sequence similarity E-values (≤ 1×10^−10^) to define neighbours (Fig 1B). Since non-allergens are more abundant than allergens in the real scenario, we designed an imbalanced dataset to simulate the case, and expected the model would be able to learn and identify allergens from proteins. For non-allergens, we collected five protein sequences not annotated as allergens from the same species of each allergen as described in the Material and Methods Training dataset section (Fig 1C).

**Figure 1.**
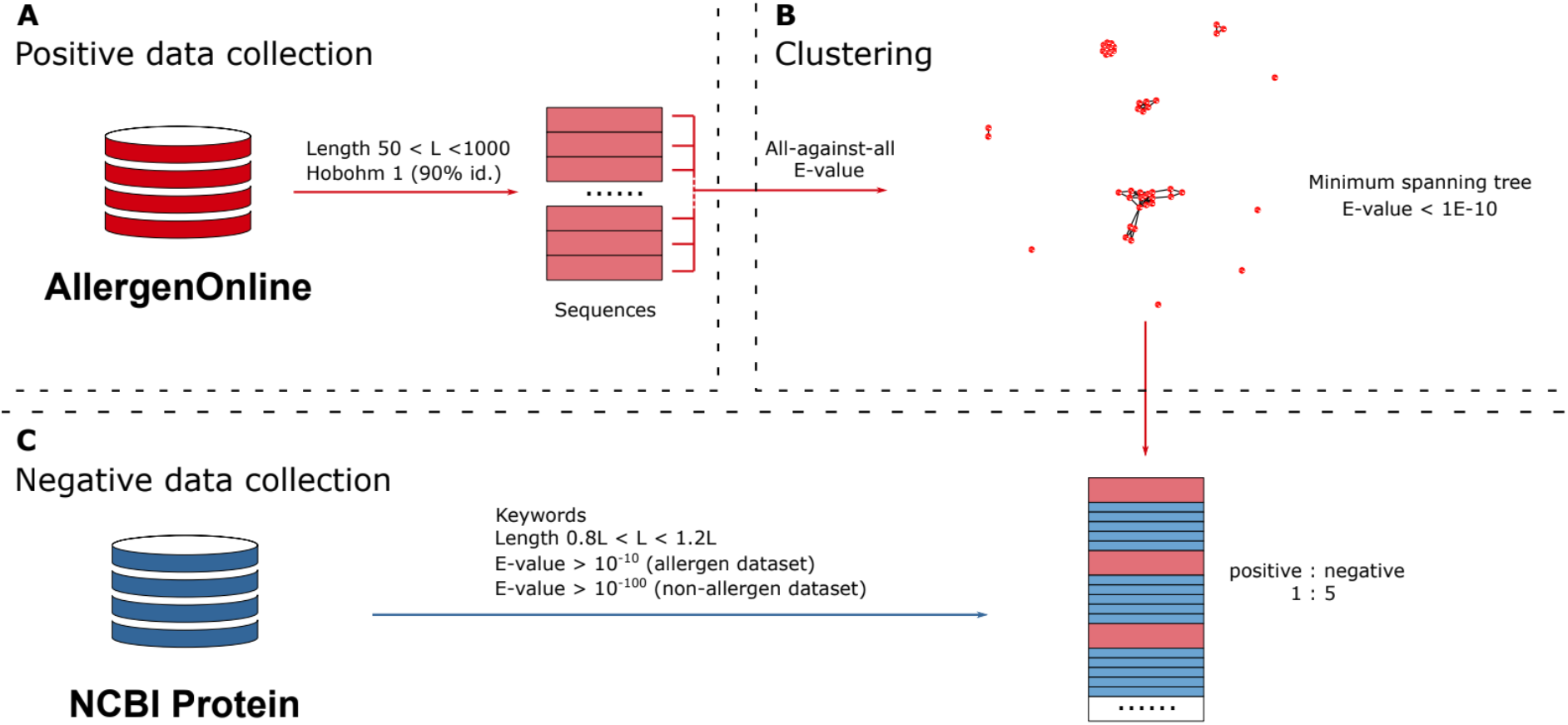
Data collection. **A**. Positive data collection. Allergens were collected from the AllergenOnline database, filtered by length, and by similarity using Hobohm 1 algorithm. **B**. Clustering. Clusters were generated using the minimum spanning tree algorithm and distances from BLAST all-against-all E-values. **C**. Negative data collection. Non-allergen sequences (blue) were collected from NCBI and added in a 5:1 ratio to positive allergens (red) searching for the same species in each allergen. When more than 5 non-allergens meet the following criteria, matching keywords, length (range 20% allergen), and similarity (E-values), a random subset of the candidates that was selected. See Materials and Methods section for details of data collection.

### MHC presentation propensity feature

In order to induce Th2 responses and IgE production, allergens are first digested by antigen-presenting cells and selected peptides are presented on MHC-II complexes. Since GWAS studies of allergic patients have shown strong association to specific MHC class II alleles, allergens contain differences on MHC-II presentable peptides to induce allergy^7–13^. NetMHCIIpan was used to predict MHC presentation potential for allergens and non-allergens on a set of selected alleles^33^. MHC-II presentation propensity was collected by adding the counts of predicted binders for a set of relevant alleles derived from genetic associations found in allergies^8,9^. In particular two features, binder 1 and binder 2, were created for grouping known positive (HLA-DRB1*04:01, HLA-DQA1*02:01-DQB1*02:02, HLA-DQA1*04:01-DQB1*03:01, HLA-DQA1*01:03-DQB1*06:01, HLA-DQA1*03:01-DQB1*03:02, HLA-

DQA1*05:01-DQB1*02:01), and the ambiguous allele (HLA-DRB1*15:01), HLA-II allergenicity associations. HLA-DRB1*15:01 was separated as it was previously proposed in some allergies as a risk factor, while in others a protective allele^11,12^.

In addition to MHC-II presentation propensity, previously described classification features for allergen/non-allergens were included. Those include amino acid compositions, physicochemical properties, structural features (alpha helix, beta sheets, and relative solvent accessibility), and evolutionary properties (see feature constructions in Materials and Methods). Feature distributions for allergen and non-allergens were illustrated by boxplots for the most significant features displaying the median, quantiles, and densities of each set of data (Fig 2). For binder 1, the t-statistic and p-value were 12.892 and 4.97×10^−34^ respectively, indicating a higher MHC-II presentation propensity on allergens compared to non-allergens. Differences in binder 2 were not statistically significant, suggesting that HLA-DRB1*15:01 presentation propensity was not discriminative for this entire dataset. However, the HLA-DRB1*15:01 showed significant presentation preference for the Der p 1 family and Bet v 1 family, compared to other allergen families, such as the Tri a 36 (wheat) and Amb a 4 (short ragweed) (Supp. Fig 2). When comparing the average residue weight, the t-test had a negative t-statistic value. These results suggested that allergens are characterised by a higher frequency of small-size amino acids. Likewise, allergens were found to be more likely hydrophilic and less hydrophobic, as previously described7, and containing less arginine and more aspartic acid. The t-statistic for alpha helix was negative, and beta sheet positive which indicated the allergens are depleted in alpha helices and enriched to beta sheets compared to non-allergens confirming previous observations^43^.

**Figure 2.**
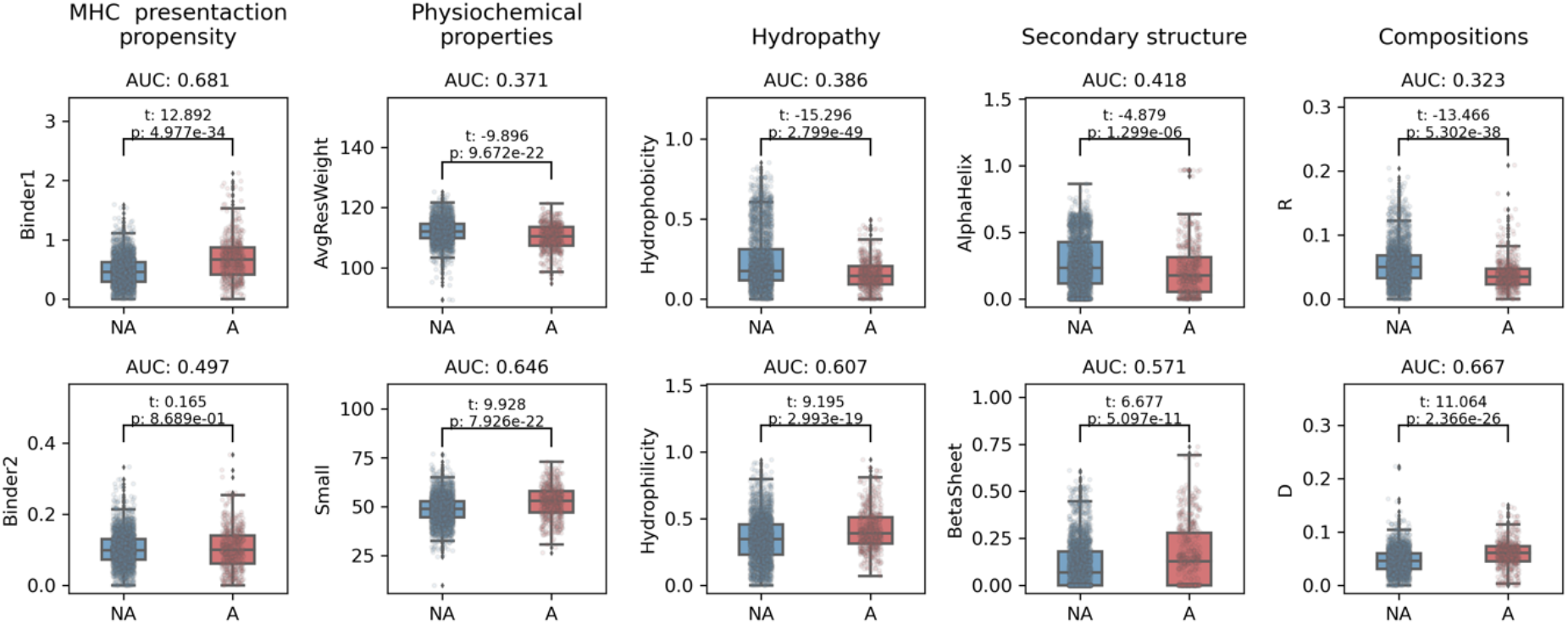
Distribution and individual-feature performances on selected features for allergens and non-allergens. T-test was used to compare the significant difference between allergens (red, A) and nonallergens (blue, NA). The AUC was calculated with feature values and the targets. “Small” corresponds to the relative abundance of small-size amino acids (A, C, D, G, N, P, S, T, V) in a given protein sequence. Likewise, D and R correspond to the frequencies of aspartic acid and arginine per protein.

We next determined the AUC of all the individual features on the training dataset. Here, AUC values below 0.5 would suggest that allergic proteins have lower feature values than non-allergic proteins. As expected, the results of these analyses aligned with the t-statistics above, i.e. allergens had more MHC-II binder 1, more small-size amino acids, fewer alpha helices, more beta sheets, more hydrophilic proteins and contained more aspartic acid and less arginine (Fig 2).

### Model development

After showing that MHC presentation propensity is a discriminatory feature for allergens (Fig 2), we sought to incorporate these features into machine learning models in order to improve the state-of-the-art allergenicity predictors. To effectively avoid overfitting, a five-fold nested cross-validation scheme was performed (Fig 3), and the best random forest hyperparameters were selected in the inner loop (Fig 3B). MHC presentation propensity was combined with different sets of features, and optimal performance on the training data was obtained with a random forest model with 60 features (60F) (Supp. Table 3). The 60F model included MHC presentation propensity, physicochemical features, amino acid compositions, auto-covariances, and structural features (Supp. Table 4, for details refer to the Material and Methods section). Finally, the partitioning from MST was repeated ten times using different random seeds and a final random forest model ensemble was generated. The ensemble model significantly outperformed most of the individual models in each random seed (Fig 3D), therefore the ensemble scheme was applied to all the models.

**Figure 3.**
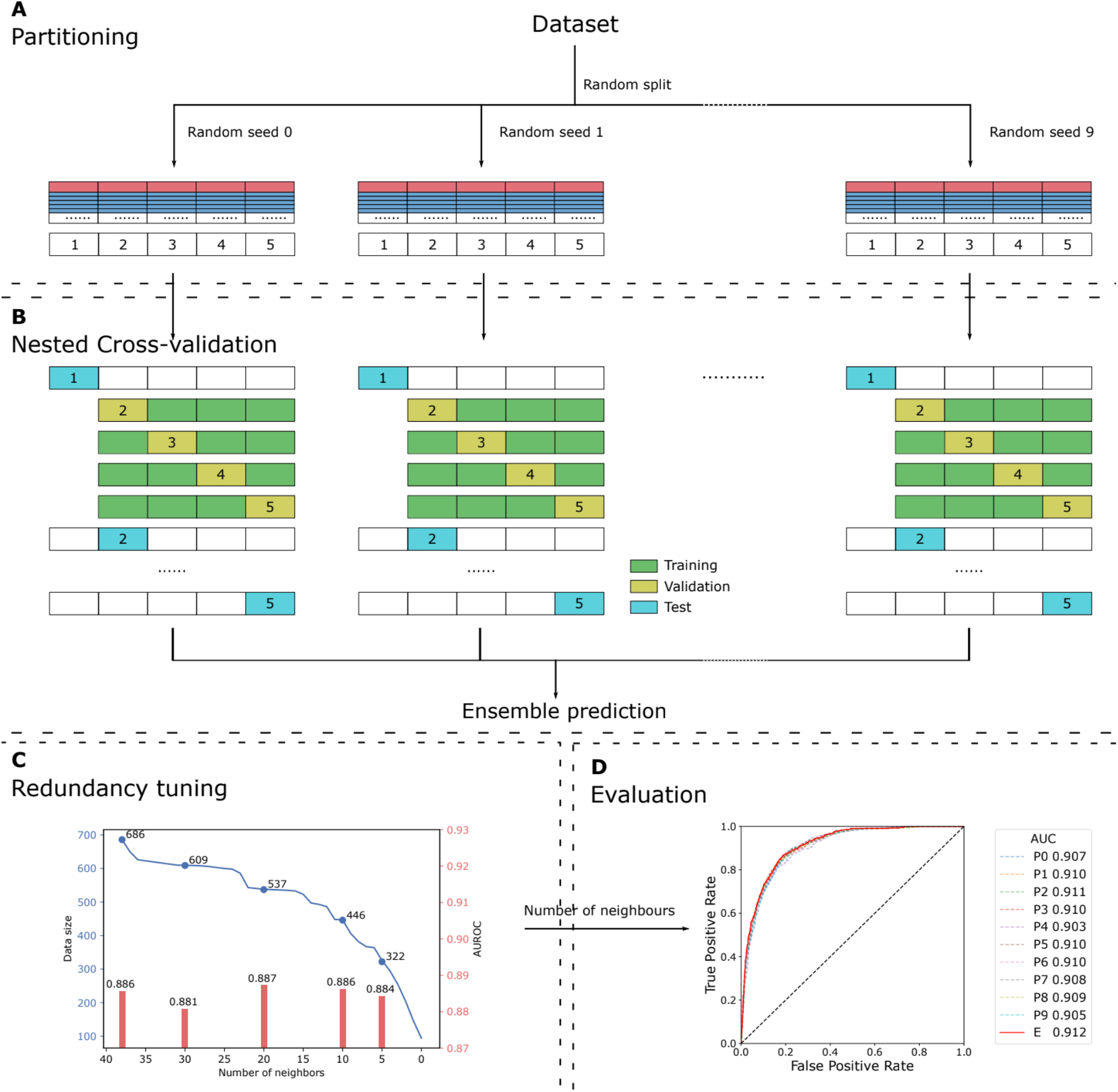
Model development pipeline of NetAllergen. **A**. Partitioning. Five partitions randomly combining the clusters obtained from minimum spanning tree (MST) clustering were distributed using ten different random seeds. Every allergen (red bar) and its five associated negatives (blue bars) share the same partition. **B**. Cross-validation and ensemble prediction. A nested cross-validation setup was used using five folds in the outer layer and four folds in the inner layer. The final ensemble prediction is the mean of the 10 model predictions. **C**. Internal redundancy tuning. The AUROC was evaluated on the minimal common subset (the dataset with a filter of 5). **D**. Evaluation. Ensemble AUC (60F) was significantly higher than the individual partitions except when comparing to P2, which was not significant (p-value 0.152).

We next assessed removing sequences with a large number of other similar sequences to further cut out redundancy in the dataset (see Materials and Methods section). Here, cross-validation predictive performance was evaluated to find the optimal filter for removing the allergens with a higher number of neighbours in the tree network after MST clustering (Fig 3C). The removal of highly connected vertices in the tree reduced the bias effect of highly similar sequences and improved the model performance. However, applying more stringent filters (removing vertices with fewer than 20 neighbours) caused a loss of informative allergen sequences as well as the performance. Hence, we applied a filter of 20 and removed the sequences that had more than 20 neighbours in the MST clusters.

### MHC presentation propensity is a predictive feature for allergenicity

Here, we picked random forest model as feature importance can be calculated as the mean decrease of Gini impurity for each specific feature in all the trees. A higher value indicates a better split of the data using that feature. We explored the contribution of the different features in the random forest model of 60F and in a reduced model of 20F (AUC=0.869) only including MHC presentation propensity (MHC), structural properties (NSP), and physicochemical features (PCP) (Supp. Table 3 and 4). MHC-II presentation propensity binder 1 was the most important feature in the simpler 20F model (Fig 4A). The frequency of small amino acids (Small), average relative solvent accessibility (AvgRSA), binder 2, frequency of acidic amino acids, and alpha helix and beta sheet were also informative features to discriminate allergens and non-allergens (above the median of all feature importance). In the 60F model, although binder 1 was no longer the most important feature, it still played an important role in the prediction (Fig 4B). In contrast, the binder 2 feature became less important in the 60F model. Further, the amino acid composition of arginine, aspartic acid, and lysine, together with the auto-covariance of PSSM on Glutamine (AC_Q) were found to be among the most discriminatory features in the 60F model.

**Figure 4.**
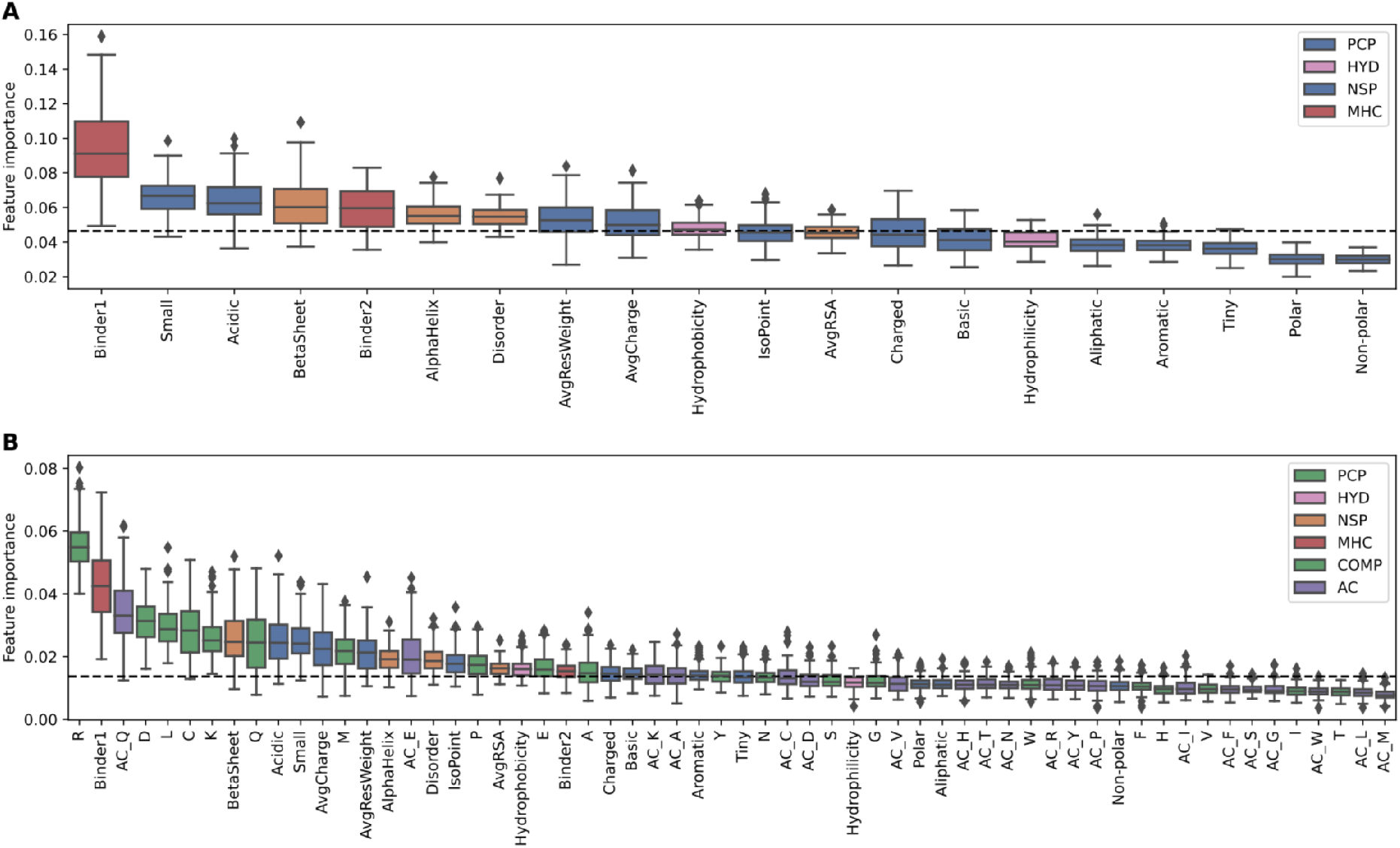
Median feature importance for random forest (RF) models. Medians are calculated from cross-validation partitions and 10 random seeds in each random forest as the mean decrease of Gini impurity for **A**. 20 features (20F) model, and **B**. 60F model. Individual features were grouped into physicochemical properties (PCP, blue), hydropathy (HYD, pink), structural information (NSP, yellow), MHC-II presentation propensity (MHC, red), amino acid compositions (COMP, green), and evolutionary information (AC, purple). Average relative solvent accessibility (AvgRSA).

To further investigate the importance of MHC-II features, models removing the binder 1 and binder 2 were trained and evaluated using cross-validation. When the MHC-II features were removed from the 60F model, AUC was reduced from 0.912 to 0.911 and AUC 0.1 from 0.512 to 0.510 (Table 1). For the 20F model however, when taking out the MHC presentation propensity features (18F), the reduction in prediction power was found to be significant for AUC and PPV). Additionally, the AUCs of 60F or 20F were both greater than AUC of any single feature (Fig 2), which indicated that the model learnt features of allergens and benefited from the combination of features. Consistently with feature importance in those models, this result indicates that the addition of amino acid compositions and autocovariances could be correlated with MHC presentation propensity. By computing the feature space, the Pearson correlation coefficient suggested that the binder1 was strongly correlated to the alanine and negatively correlated to cysteine (Supp. Fig 3). These features contributed to the prediction when the MHC presentation propensity was absent, which led to no significant difference after the removal of binder features.

**Table 1.**
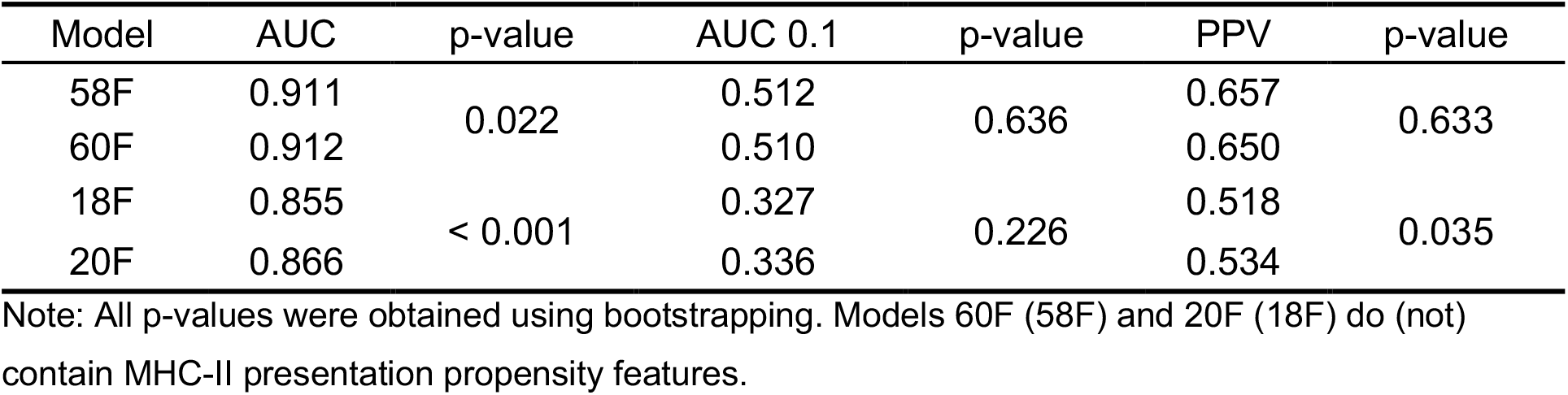
Cross-validation performance of models with and without MHC-II features.

### Model comparison

Sequence similarity to previously known allergens is the main driver of previously developed allergenicity predictors^17,44^. Therefore, we constructed a sequence similarity baseline model using BLAST. In the baseline model, the predicted value for each potential allergen from the evaluation datasets was taken from the best hit of BLAST against the training dataset. This baseline model was compared to the random forest model (60F) on the AlgPred 2 evaluation dataset (see Materials and Methods). Overall, the baseline model had a higher performance (AUC of 0.923) on the full evaluation dataset compared to the random forest of 60F (AUC 0.886, Supp. Table 5). This exceptional performance of the baseline model clearly suggests the high similarity between the allergens in the evaluation dataset and those in the training dataset. However, it is expected that the performance of the baseline model will decrease on unknown or remotely related allergen sequences. To quantify how model-performance correlates with sequence similarity to the training dataset, we sorted the sequences in the evaluation dataset from high to low similarity (negative log E-value, i.e., the predicted values of baseline model). We observed that the baseline model performed slightly better than our 60F model for highly similar query entries to the training data, but that dropped dramatically when sequence similarity decreased (Fig 5). On the contrary, the 60F model demonstrated a more robust AUC performance on all the dataset and outperformed the baseline model on proteins that were less similar (E-value > 1×10^−51^) to sequences in the allergen database (Fig 5A). The same trend was observed for AUC 0.1 (Fig 5B).

**Figure 5.**
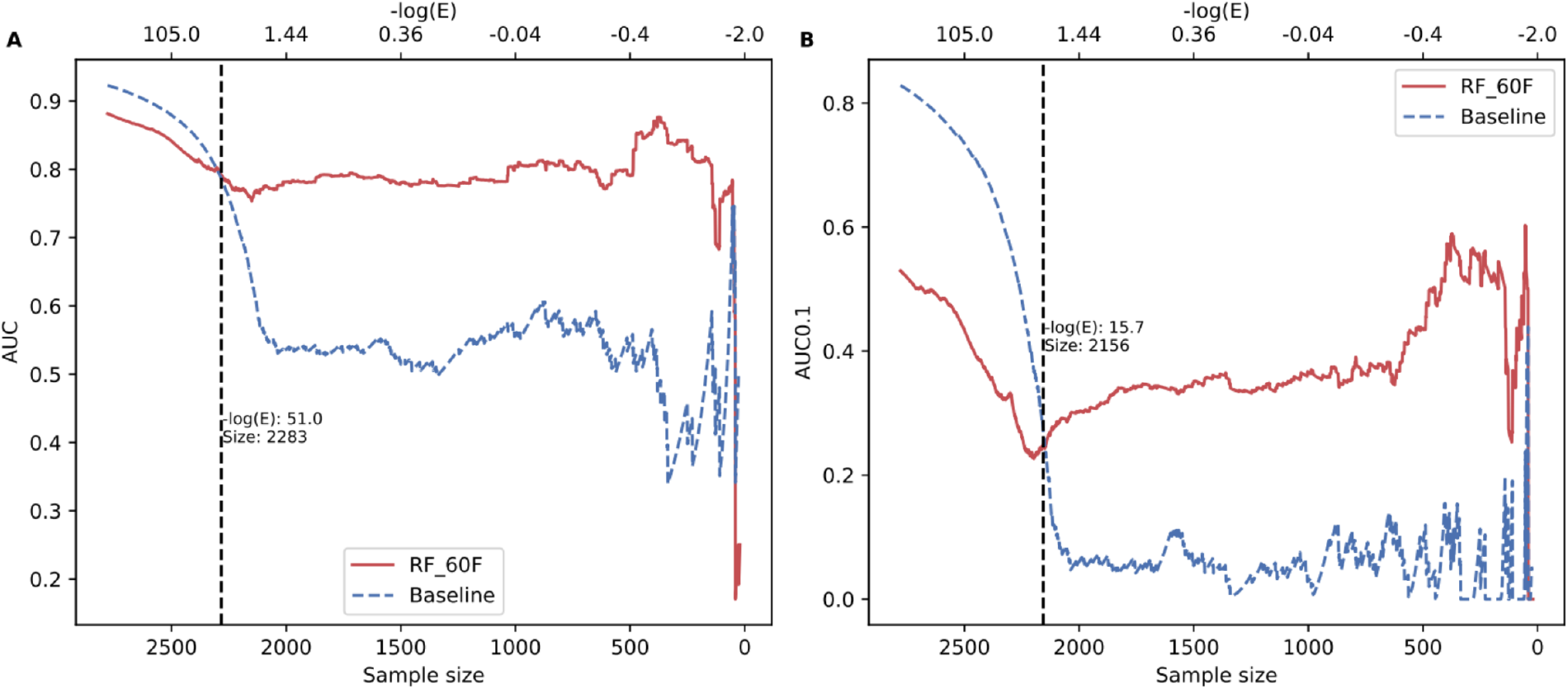
Model performance for variable datasets with decreasing similarity thresholds on AlgPred2 evaluation dataset. The sequences were sorted by similarities against allergens in the training dataset with a descending order. The vertical line indicates the changes of performance and split the curves into areas of higher and lower similarities. For the rightmost curve, the number of sequences reduced as the decreasing similarity, which led to that the evaluation was less robust because of noise. **A**. AUC. **B**. AUC 0.1.

In order to compare to state of the art allergenicity predictors, we next benchmarked the random forest model against AlgPred 2^27^. Since AlgPred 2 was available in a re-trained format using the evaluation dataset used in their publication, we collected a second and independent evaluation dataset from other allergen databases (detailed information in Methods). As in the earlier benchmark, the new independent validation set was ranked in the same method with E-values against the allergens in the combined datasets of the cross-validation and AlgPred 2. Both AUC and AUC 0.1 performances were assessed in the identical approach and behaved similarly as before, showing a higher performance of AlgPred 2 and baseline model for the highly similar sequences (Fig 6). In a similar fashion, both AlgPred 2 and BLAST performances dropped below that of the 60F model when similarity to training-allergens E-values were higher than 1×10^−59^. This result suggests the 60F model had superior predictive power on novel sequences which were not included in current allergen databases.

**Figure 6.**
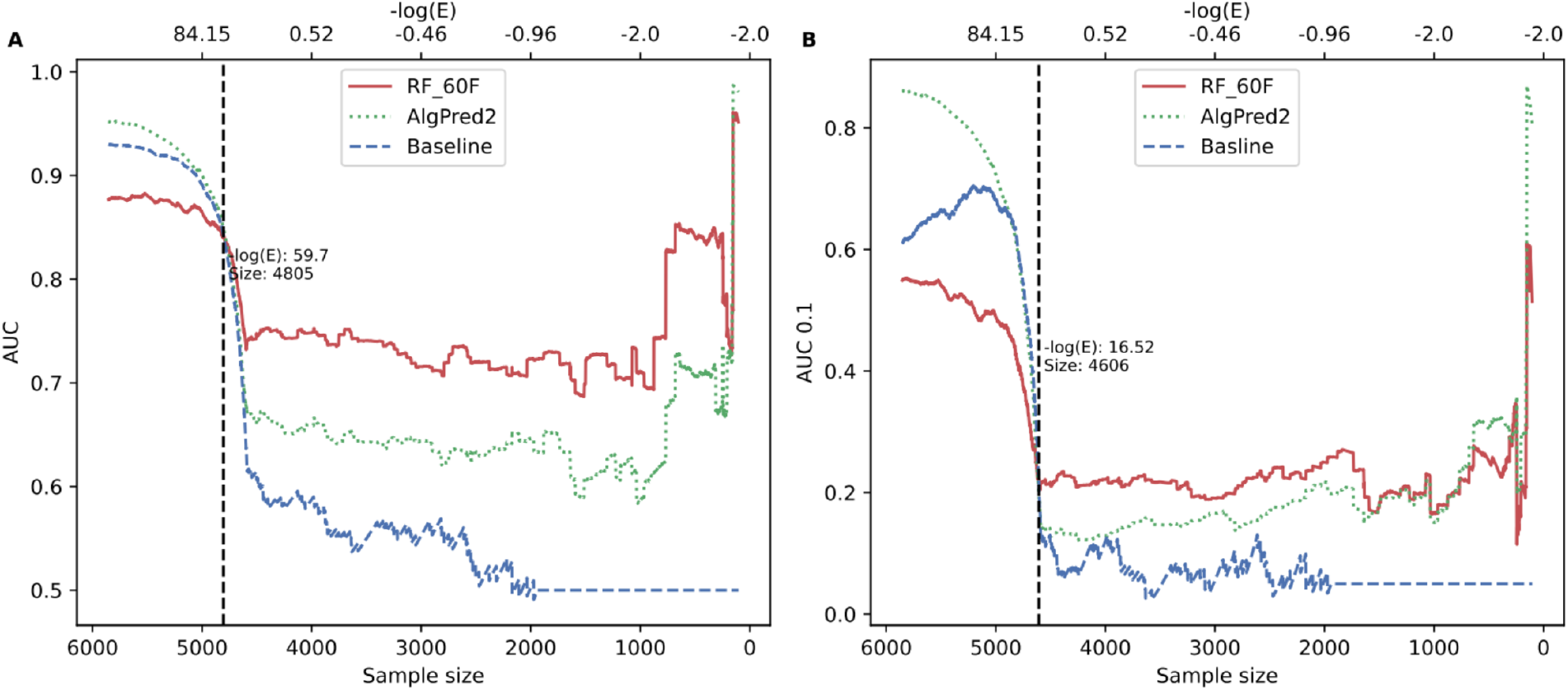
Model performance with different similarity thresholds on the new evaluation dataset. The similarities were represented by -log(E-value) which were obtained from the baseline model (BLAST). The searching database consisted of the positive sequences from our and AlgPred2 dataset. The noise in decreasing data size led to the less robust curves at the rightmost part. **A**. AUC. **B**. AUC0.1.

### Web server

The 60F random forest model was deployed as a web server. The allergenicity predictor is available at https://services.healthtech.dtu.dk/service.php?NetAllergen-1.0. The program accepts a FASTA file of protein sequences and provides a prediction result table (Fig 7).

**Figure 7.**
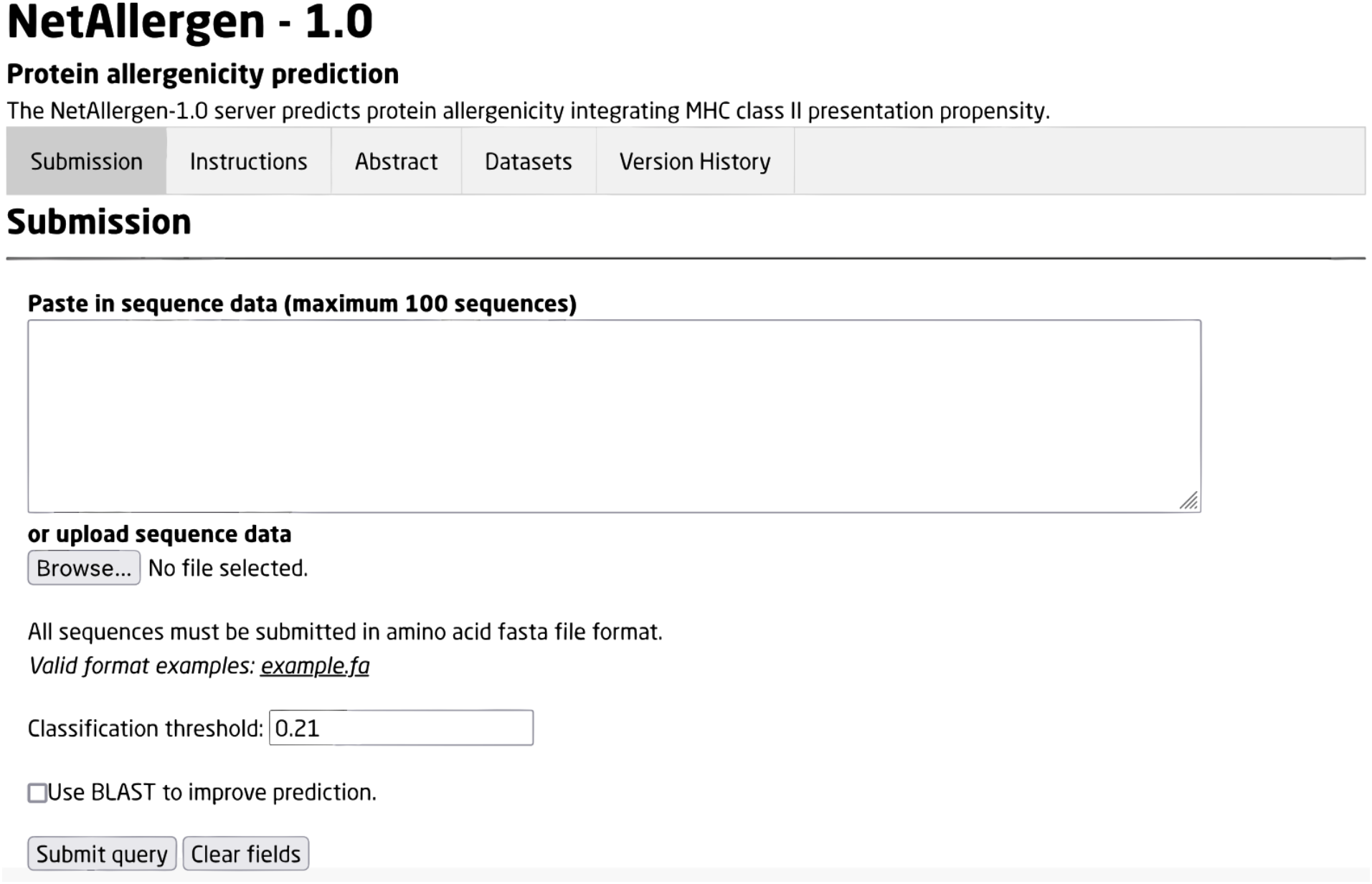
Web service of NetAllergen-1.0.

## DISCUSSION

Understanding the nature of allergenicity is of high importance as it contributes to the development of future diagnostic tools and therapeutic strategies. Previous allergen predictors have focused on answering the question of what makes an allergen. By introducing HLA presentation propensity, we proposed a first attempt to answering the question of why only some individuals develop allergies towards ubiquitous proteins. Here, we have developed a new allergen predictor called NetAllergen which incorporates a novel significant feature regarding MHC class II presentation on a set of particular susceptible alleles, suggesting allergenicity may relate to presentation on particular subset of HLA class II alleles. NetAllergen was trained on a carefully curated dataset where redundant allergen sequences were removed. Due to the accessibility of both program/web service and training dataset, AlgPred 2 was the only accessible tool used for benchmark. NetAllergen was demonstrated to perform better than a state-of-the-art predictor AlgPred 2, and a baseline model when sequence similarity towards known allergens is reduced.

Although it has long been recognised that allergy has a strong HLA association component^45^ still very little is known about the mechanisms that trigger allergy. Recently, large genome-wide association studies pointed to conserved specific amino acid positions in the HLA-II binding groove determining protective or risk variants for certain allergies^7,8^. This observation suggests that MHC-II presentation in those alleles might have an important role in allergy. Consequently, we have hypothesised that allergens might have a high quantity of presentation motifs towards those specific alleles compared to non-allergens. Here, we showed that this is indeed the case. HLA allele presentation propensity in binder 1 was one of the strongly discriminant features for allergens (AUC=0.68). In addition, incorporation of this novel HLA presentation propensity feature with other structural features, amino acid compositions, etc, into random forest models achieved good performance for allergenicity prediction.

The advantage of random forest compared to other more complex models in machine learning is its ease of interpretability. Therefore, we have picked this simple model to better analyse feature importance. Once more, MHC presentation propensity binder 1 was found to be one of the most important features when analysing which features best split the data in the random forest. Moreover, removing MHC class II presentation features from the model (20F) significantly reduced performance compared to a model including those features. This suggested that MHC-II presentation is an informative classification allergen feature. However, when incorporating additional features such as amino acid compositions, and evolutionary information of the amino acids, those improvements were only moderate. In the 60F models, Arginine frequency was the most important feature followed by MHC-II presentation binder 1, auto-covariance of glutamine (AC_Q), and amino acid compositions of aspartic acid, leucine, cysteine and lysine. While MHC-II presentation features are based on the count of presentation motifs to specific MHC-II alleles, the amino acid compositions count the percentage of amino acids in a protein sequence. They both capture important amino acids information in two distinct approaches, explaining why the 60F model did not show significant improvements against the 58F model.

In addition to the MHC-II binders, the average residue weight was negatively correlated to allergenicity and small amino acids composition (A, C, D, G, N, P, S, T, V) had positive correlation. In particular, threonine and valine are branched on the beta-carbon and thus are more often found on beta strands than in alpha helices. On the other hand, proline is disfavoured on alpha helices as its amino ring is not able to adopt normal helical main chain conformations^46^. These findings align with previous observations that allergens have a higher abundance of beta sheets and less alpha helices compared to non-allergens^43^.

Compositions related to Arginine and Aspartic acid tended to have respectively a negative and positive correlation to allergy, in agreement with more acid amino acid compositions. Auto-covariance (AC) represents the average covariance between the sequence and itself. Hence, the coefficients of AC_Q indicated that Glutamine might be present in a large number of repetitions in the sequence. This information suggested by AC of Glutamine was similar to the Q-repeats features used in AllerCatPro, thus aligning with previous findings for allergens^24^.

Many important applications in bioinformatics, including sequence alignment and machine learning algorithms, employ sequence weighting schemes to mitigate the effects of non-independent homologous sequences over-represented in a dataset^47,48^. In particular, allergen datasets are highly redundant as many homologous proteins are discovered in closely related species. Moreover, strict sequence similarity guidelines proposed by the FAO/WHO to assess allergenicity in-silico may have favoured the discovery of iso-allergens. As part of the development of NetAllergen, we have proposed a novel pipeline to deal with highly redundant datasets and avoid data leakage issues. Here, we have showcased how this pipeline can be applied to real-life datasets aiding the development of data-driven prediction methods. We believe this proposed pipeline to be broadly applicable and expect it to be of general use for the community.

## DATA AVAILABILITY

The data used in this manuscript is available at the web server link; https://services.healthtech.dtu.dk/service.php?NetAllergen-1.0.

## AUTHOR CONTRIBUTIONS

YL collected the data, developed the pipeline, trained the models, analysed and interpreted the data; PWS created the web server; MN analysed and interpreted the data, provided critical revision of the article; and CB analysed and interpreted the data, provided critical revision of the article, and conceived the work. All authors wrote the paper and approved the final version to be published.

## CONFLICT OF INTEREST

All authors declare no competing interest.

## Notes

### Competing Interest Statement

The authors have declared no competing interest.

### Summary of Updates

updated version

