## Supplementary Material for "NetAllergen, a random forest model integrating MHC-II presentation propensity for improved allergenicity prediction"

**Supp. Table 1. Hyperparameters tuning in random forest**

| Hyperparameters | Range |
| --- | --- |
| Number of estimators | 40–200 with a step of 20 |
| Depth of trees | 4–20 with a step of 2 |
| Minimum samples for split | 2, 4, 8 |
| Minimum samples on a leaf | 1, 2, 4 |
| Impurity criteria | gini, entropy |

**Supp. Table 2. The composition of the evaluation dataset**

| Database | Positive data size | Negative data size | Data size |
| --- | --- | --- | --- |
| Allergen Nomenclature | 106 | 487 | 593 |
| AllerBase | 136 | 682 | 818 |
| COMPARE | 27 | 131 | 158 |
| Allergome | 767 | 3,514 | 4281 |
| SDAP | 1 | 5 | 6 |
| Total | 1,037 | 4,819 | 5,856 |

Note: The allergen protein sequences that were identical to allergens already present in the training dataset were excluded. The non-allergen sequences were collected following the same criteria as for the training dataset (detailed in Material and Methods section).

**Supp. Table 3. Cross-validation model performance with incremental feature spaces**

| Model | AUC | AUC 0.1 | PPV |
|---|---|---|---|
| 20F | 0.869 | 0.348 | 0.544 |
| 40F | 0.908 | 0.494 | 0.644 |
| 60F | 0.913 | 0.509 | 0.646 |

Note: The random forest feature spaces were divided into three different subspaces: 20F, 40F and 60F. The 20F model included physicochemical properties, local structural features, and MHC class II presentation propensity. The 40F model included the 20F features + 20 amino acid compositions. The 60F model included 40F + 20 features for evolutionary information for each amino acid (auto-covariance). Detailed information about the feature construction is explained in Material and Methods section.

**Supp. Table 4. Feature compositions in the random forest models**

| Model | MHC presentation propensity (2) | Physicochemical properties (12) | Hydropathy (2) | Secondary structure (4) | Composition (20) | Evolutionary information (20) |
|---|---|---|---|---|---|---|
| 18F | – | + | + | + | – | – |
| 20F | + | + | + | + | – | – |
| 40F | + | + | + | + | + | – |
| 56F | + | + | + | – | + | + |
| 58F | – | + | + | + | + | + |
| 60F | + | + | + | + | + | + |

Note: The name of the model represents the number of features included in the model. “+” indicates that the features were included in the current model, while “–” indicates non-included features.

**Supp. Table 5. AlgPred2 validation dataset benchmark**

| Model | AUC | AUC 0.1 | PPV |
|---|---|---|---|
| RF_60F | 0.886 | 0.551 | 0.712 |
| Baseline | 0.923 | 0.829 | 0.848 |

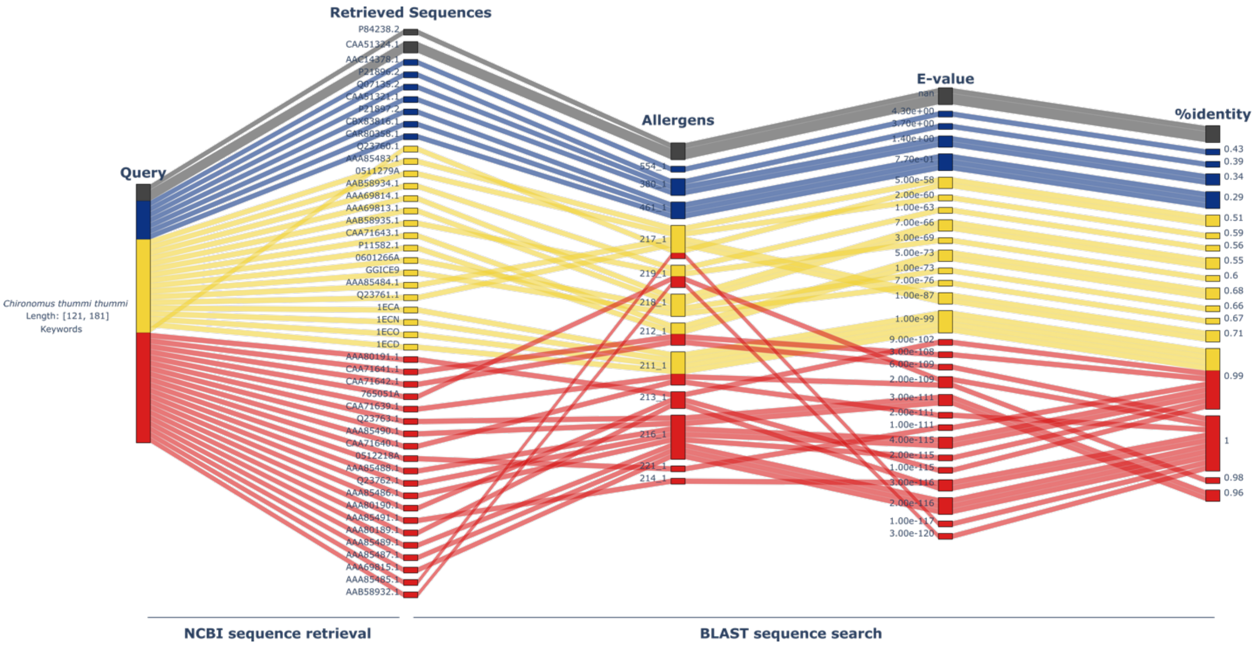


**Supp. Figure 1.** Sequence alignment of non-allergen candidates against the allergen dataset for negative dataset collection. Take an allergen (from *Chironomus thummi thummi*) as a query, 47 retrieved sequences were downloaded by searching the NCBI protein database with the species name, length filter and keywords (see Materials and Methods). Red line indicates the non-allergen candidates are highly similar (< $1E^{-100}$) to allergens in our dataset. The candidates with an E-value between $1E^{-100}$ and $1E^{-10}$ are labelled in yellow. Blue line means the candidate sequences share low similarity with allergens. Those not found in BLAST are grey. Random selection of candidates without filters is likely to have mislabel problems in training.

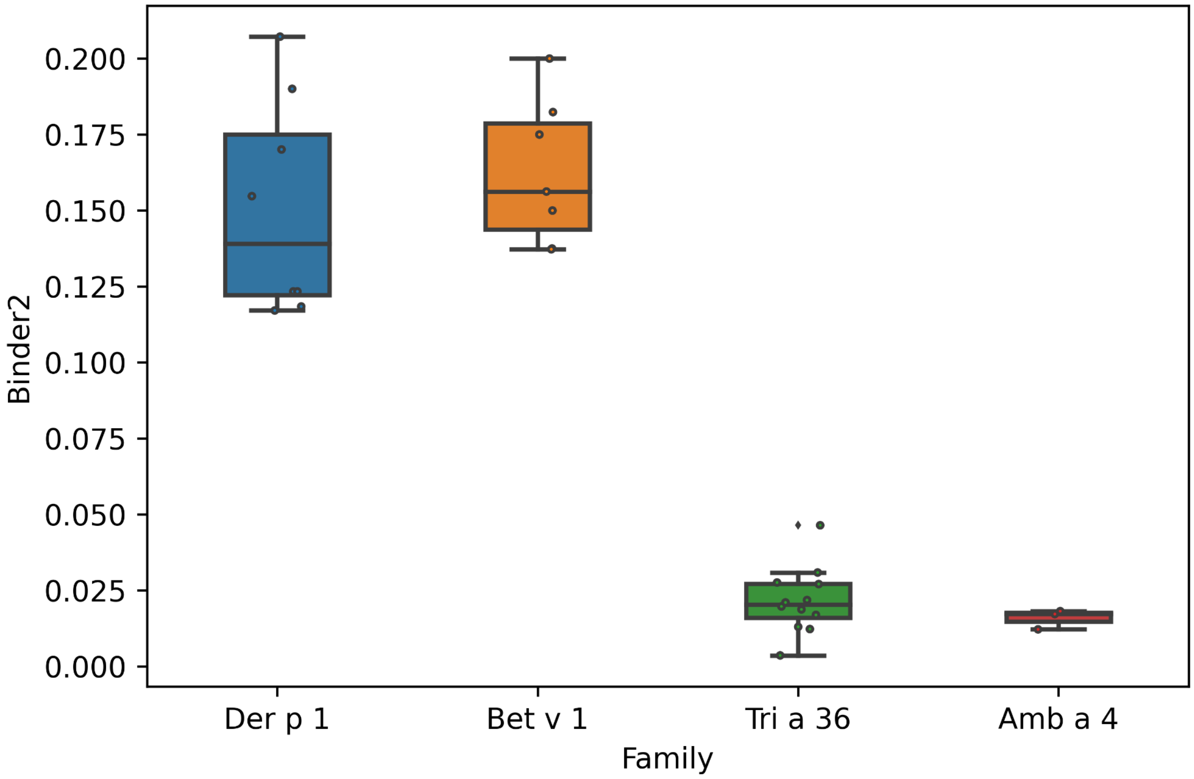


**Supp. Figure 2.** The distribution of binder2 feature (DRB1*15:01) of four types of allergen families. The Der p 1, Bet v 1 families have higher propensity of being presented by DRB1*15:01 MHC molecules, which aligns with previous studies about association between DRB1*15:01 and allergy.

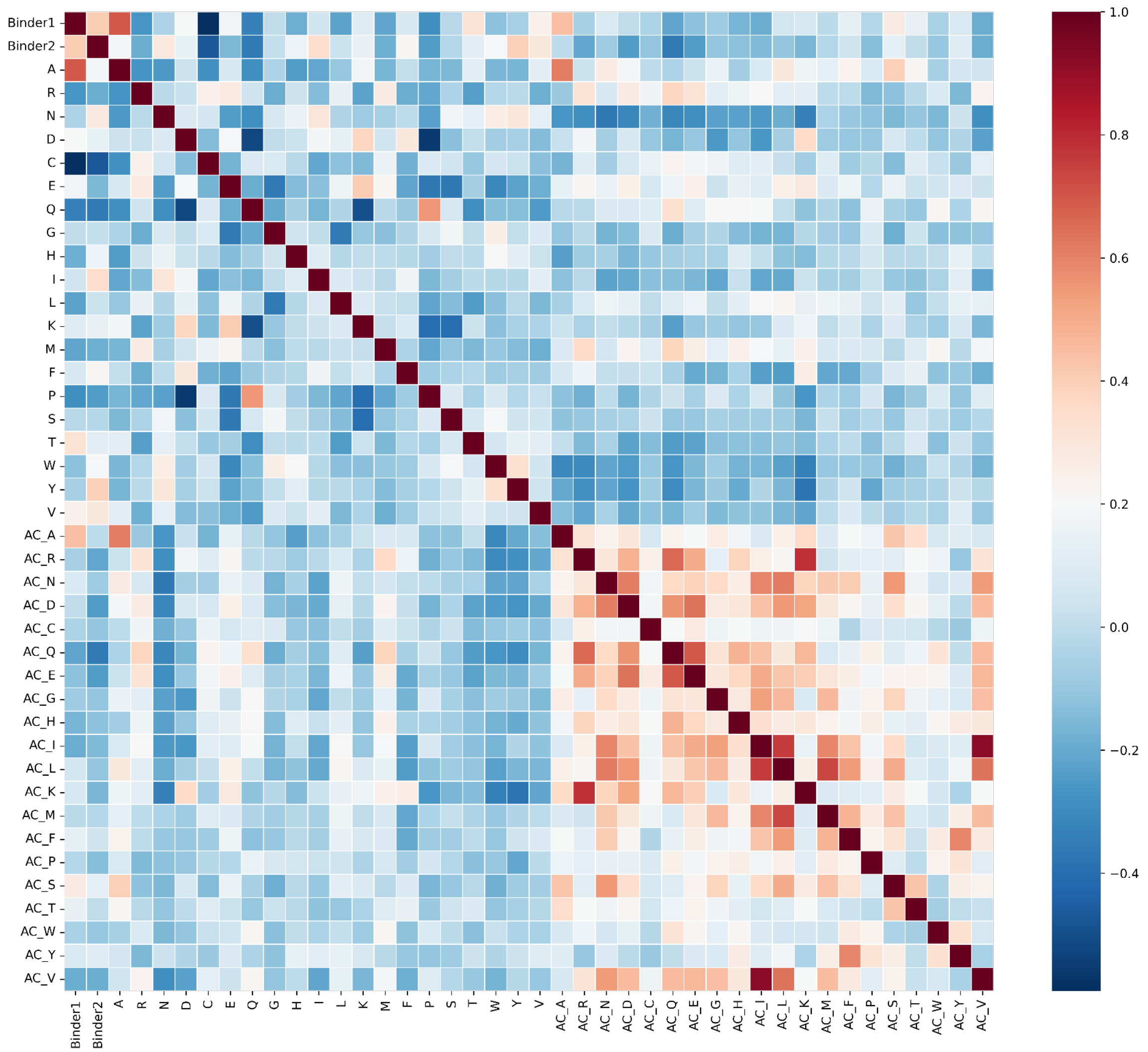


**Supp. Figure 3.** Pearson correlation coefficient of MHC II presentation features (binder1, binder2), amino acid compositions (A, R, N, etc.) and autocovariances (AC_A, AC_R, AC_N, etc.) for allergens.
